# Cannabis use is associated with increased levels of soluble gp130 in schizophrenia but not in bipolar disorder

**DOI:** 10.1101/806927

**Authors:** Attila Szabo, Ibrahim A. Akkouh, Thor Ueland, Trine Vik Lagerberg, Ingrid Dieset, Thomas Bjella, Pål Aukrust, Stephanie Le Hellard, Anne-Kristin Stavrum, Ingrid Melle, Ole A. Andreassen, Srdjan Djurovic

## Abstract

The complex effects of plant cannabinoids on human physiology is not yet fully understood, but include a wide spectrum of effects on immune modulation. The immune system and its inflammatory effector pathways are recently emerging as possible causative factors in psychotic disorders. The present study aimed to investigate whether self-administered cannabis use was associated with changes in circulating immune and neuroendocrine markers in schizophrenia (SCZ, n=401) and bipolar disorder patients (BD, n=242). A screening of 13 plasma markers reflecting different inflammatory pathways was performed in SCZ and BD patients after subdividing each group into cannabis user and non-user subgroups. We found that i) soluble gp130 (sgp130) concentrations were significantly elevated among cannabis users in the SCZ group (p=0.002) after multiple testing correction, but not in BD. ii) Nominally significant differences were observed in the levels of IL-1RA (p=0.0059), YKL40 (p=0.0069), CatS (p=0.013), sTNFR1 (p=0.031), and BDNF (p=0.020), where these factors exhibited higher plasma levels in cannabis user SCZ patients than in non-users. iii) These differences in systemic levels were not reflected by altered mRNA expression of genes encoding sgp130, IL-1RA, YKL40, CatS, sTNFR1, and BDNF in whole blood. In sum, our results show that cannabis self-administration is associated with markedly higher sgp130 levels in SCZ, but not in BD, and that this phenomenon is independent of the modulation of peripheral immune cells. These findings warrant further investigation into the potential neuroimmune, anti-inflammatory, and biobehavioral-cognitive effects of cannabis use in SCZ.

## BACKGROUND

The psychoactive and medicinal properties of the *Cannabis indica* plant, a species of the genus *Cannabis* of Cannabaceae family, have been recognized for millennia for its therapeutic value in inflammation, pain, and rheumatic diseases (Russo, 2007). The biologically active cannabinoids can be classified into two major groups based on their psychotropic effect. The psychoactive delta-9 tetrahydrocannabinol (Δ9-THC or THC) and the non-psychoactive cannabidiol (CBD) are the two most widely studied cannabinoids with potential therapeutic value. However, beyond their effects on immunity and inflammation, cannabinoids have also been shown to modulate neuronal development, as well as to influence the general cytoarchitecture of the brain by regulating axon and dendrite growth, synaptic dynamics, and pruning (Njoo *et al*., 2015). This latter phenomenon has recently been linked to their capacity to increase the levels of neurotrophins in the mammalian CNS, such as brain-derived neurotrophic factor (BDNF) (Suliman *et al*., 2018; Sales *et al*., 2018). It has also been associated with potential adverse effects with regards to cognition, behavior, respiratory, and cardiovascular disorders (Curran *et al*., 2016; Cohen *et al*., 2019).

Schizophrenia (SCZ) and bipolar disorder (BD) are severe mental illnesses which pose a substantial burden on the global community by greatly affecting the mortality, life quality, and costs of patient care (Correll *et al*., 2015). Although both have a strong genetic component, details of their pathophysiology and etiology is not yet known and the understanding of the underlying disease mechanisms will expectedly be crucial for developing new and effective therapies (Editorial, 2010). The immune system and its inflammatory effector pathways and elements are recently emerging as possible contributing factors in psychotic disorders (Network and Pathway Analysis Subgroup of Psychiatric Genomics Consortium, 2015). Inflammatory cytokines may alter brain functions in multiple ways, and may pass the blood-brain-barrier through leaky regions or via passive and active transport mechanisms. Plant cannabinoids have been shown to exert a wide spectrum of effects on human immunity ranging from anti-inflammation to neuroimmune modulation (Olah *et al*., 2017). THC and CBD are known to have disparate influences on SCZ and BD symptoms, and probably on their pathophysiology as well.

Despite the intensive research efforts in the past decades, the complex effects of plant cannabinoids on human physiology, including effects on the immune system, are not yet fully understood, partly due to law regulations of plant-based exocannabinoids in many countries. Since medical cannabis has already entered mainstream medicine, it is necessary for the medical community to make efforts assessing the possible risks and benefits of plant cannabinoids in order to provide reliable and competent clinical care. In this study, we aimed to investigate the effects of cannabis consumption (users versus nonusers) on circulating inflammatory, immune, and neuroendocrine markers in SCZ and BD patients. The following markers were selected because they reflect different parts of the inflammatory network: interleukin 1 receptor antagonist (IL-1RA), soluble glycoprotein 130 (sgp130), and soluble tumor necrosis factor receptor 1 (sTNFR1), which are general markers of inflammation; myeloperoxidase (MPO) and cathepsin S (CatS), which are markers of neutrophil activation; YKL40, which is a marker of monocyte/macrophage activation; chemokine C-X-C motif ligand 16 (CXCL16), pentraxin 3 (PTX3), osteoprotegerin (OPG), von Willebrand factor (vWF), and activated cell adhesion molecule (ALCAM), which are markers of vascular inflammation; Parkinson disease protein 7 (PARK7) and brain-derived neurotrophic factor (BDNF), which are markers of neuroendocrine activation in various neuropsychiatric disorders. We and others have previously shown that some of these markers are dysregulated in psychosis (e.g. IL-1Ra, OPG, sTNFR1, vWf and sgp130), while some are related to neuroinflammation (e.g. CXCL16 and ALCAM) and can be modulated by cannabinoids (Wojkowska *et al*., 2014; Curran *et al*., 2016; Helle *et al*., 2016; Sales *et al*., 2018; Cohen *et al*., 2019; Morch *et al*., 2019). Here we report the modulatory potential of cannabis on circulating immune and neuroendocrine markers in SCZ in a disease-specific manner.

## METHODS

### Setting and participants

The current study is part of the ongoing Thematically Organized Psychosis (TOP) study, which includes participants from hospitals in the Oslo region, Trondheim, and Southeast regional hospitals in Norway. Information about recruitment procedures, inclusion and exclusion criteria, and clinical assessments for the TOP study as a whole have been described in detail in previous reports (Simonsen *et al*., 2011; Morch *et al*., 2019). All participants have given written consent and the study was approved by the Norwegian Scientific Ethical Committees and the Norwegian Data Protection Agency. The authors assert that all procedures contributing to this work comply with the ethical standards of relevant guidelines and regulations.

### Sample characteristics

The primary study sample of the present work consisted of 242 bipolar disorder patients (BD; 164 type I, 66 type II, 12 not otherwise specified) and 401 schizophrenia spectrum patients (SCZ; 296 schizophrenia, 77 schizoaffective, 28 schizophreniform). All participants underwent a clinical examination that included diagnostic interviews based on Structured Clinical Interview in DSM-IV axis I Disorders (SCID-1) and structured assessments of clinical symptoms, BMI, use of psychotropic medication, smoking habits, alcohol consumption, and illicit substance use. Diagnostic evaluation was performed by trained psychologists and psychiatrists, all of whom participated regularly in diagnostic meetings supervised by professors in psychiatry. The main inclusion criteria were confirmed diagnosis of either SCZ spectrum disorder or BD according to the Diagnostic and Statistical manual of Mental Disorders (DSM)-IV, age between 18-65, and ability to give informed written consent. The main exclusion criteria were clinically significant brain injury, neurological disorder, and any substance use other than cannabis within the last six months prior to blood sampling. Patients with ongoing infection, autoimmune disorders or cancer were also excluded. Cannabis was self-administered in the user cohorts and “cannabis use” was defined as any use in the last six months prior to blood sampling. Samples were subjected to high sensitivity plasma EIA and Illumina microarray analyses. Substance use was documented through interviews, urine samples, the Clinical Drug Use Scale (DUS) and the Clinical Alcohol Use Scale (AUS) (Drake *et al*., 1990; Drake *et al*., 1996). Substance use disorders were diagnosed using the SCID-E module. All accessible information in each case was examined to avoid false positive and false negative substance users. The study sample also included 613 healthy controls (284 females, 329 males, mean age 33.4). Since sufficient data on illegal drug use was not available for the controls, they were not included in the statistical analyses, but they were used to determine the average plasma concentrations in healthy subjects for the examined biomarkers.

### Measurement of inflammatory and neuroendocrine markers in plasma

Blood samples from both disease cohorts were obtained at Oslo University Hospital (OUH) following the same standardized procedures in accordance with the EUSTAR guidelines on biobanking (Beyer *et al*., 2011). Peripheral blood samples were collected in EDTA tubes and were centrifuged at >2100 g at room temperature within 30 minutes. Plasma aliquots were then stored at −80°C until assayed. Circulating YKL40, CatS, sTNFR1, BDNF, sgp130, IL-1RA, Alcam, MPO, CXCL16, Park7, vWF, OPG, and PTX3 levels were analyzed in duplicate using commercially available antibodies (all from R&D Systems, Minneapolis, MN, USA; except IL-1Ra, Peprotech, RockyHill, NJ, USA) in a 384 format using a combination of a SELMA (Jena, Germany) pipetting robot and a BioTek (Winooski, VT, USA) dispenser/washer. Absorption was read at 450 nm with wavelength correction set to 540 nm using an ELISA plate reader (Bio-Rad, Hercules, CA, USA). Intra- and inter-assay coefficients of variation were <10% for all EIAs.

### RNA expression analysis

Blood samples for mRNA expression measurements were collected in Tempus Blood RNA Tubes (Life Technologies Corporation). Total RNA was extracted with the TEMPUS 12-Port RNA Isolation Kit (Applied Biosystems) and ABI PRISM 6100 Nucleic Acid PrepStation (Applied Biosystems) according to the manufacturer’s protocol. Gene expression analysis of immune-related genes was performed with Illumina HumanHT-12 v4 Expression BeadChip (Illumina, Inc.). Multidimensional scaling and hierarchical clustering were used for regular quality control, including sample quality measurements and removal of outliers, as well as removal of multiple batch effects. This was followed by log2-transformation. If more than one probe for a single gene were available, only the constitutive probe targeting all gene transcripts was used. In total, 6 mRNA probes targeting six cannabis-associated immune marker genes were examined. More details on microarray pre-processing and quality control have previously been reported (Akkouh *et al*., 2018).

### Statistical analyses

Within each diagnostic category (BD and SCZ), patients were divided into two groups based on whether they had used any cannabis within the last six months prior to blood sampling (users) or not (non-users). To identify significant differences between users and non-users in clinical metrics of interest, Student’s two-sided t-test was used for numerical variables (the Wilcoxon rank sum test was used for non-parametric variables) and Pearson’s Chi-squared test was used for categorical variables. To investigate whether plasma concentrations of inflammatory and neuroendocrine biomarkers were related to cannabis use within each diagnostic group, separate multiple linear regression analyses were performed for BD and SCZ. The predictor variable was cannabis status (non-user vs. user) and the response variable was circulating immune marker concentrations. All analyses were adjusted for sex, age, BMI, smoking status (yes/no), alcohol consumption, and the use of four different types of medication (antipsychotics, anticonvulsants, antidepressants, and lithium). Medication doses were standardized prior to analysis by converting them to Defined Daily Dosages (DDD) in accordance with the guidelines from the World Health Organization Collaborating Center for Drug Statistics Methodology (https://www.whocc.no/). The Bonferroni procedure was used to adjust the p-values for the number of statistical tests performed. Because of the unbalanced design of our datasets, we log2-transformed any immunological marker with severely non-normal residuals or unequal variances between groups. To examine whether cannabis status was associated with gene expression levels of the significant inflammatory markers, a separate but partially overlapping sample of 249 BD patients (180 type I, 54 type II, 15 not otherwise specified) and 383 SCZ patients (270 schizophrenia, 78 schizoaffective, 35 schizophreniform) was used (Supplementary Figure S3, Supplementary Table S1). The same statistical procedures as described above were followed. All analyses were performed in R 3.4.1.

## RESULTS

Demographic and clinical variables are summarized in Table 1. In both diagnostic groups, age was strongly associated with cannabis use, and cannabis users were on average 6.1 years younger than non-users. Among SCZ patients, strong associations were also found between cannabis use and BMI (p=6.54e-5), tobacco smoking (p=3.93e-9), alcohol consumption (p=6.91e-7), and age of onset of psychotic symptoms (1.34e-4). Interestingly, BD patients who smoked cannabis experienced less negative symptoms than patients without recent cannabis use (p=0.009). In the SCZ group, cannabis use was associated with an increased risk of psychotic symptoms in total (p=0.039), which is in agreement with the recent literature (Helle *et al*., 2016; Cohen *et al*., 2019). There were no significant differences in the use of antipsychotics, antiepileptics, or lithium in any of the disease groups, but SCZ patients who smoked cannabis used less antidepressants than non-smokers (p=0.027).

**Table 1.** Demographic and clinical characteristics of schizophrenia (SCZ) and bipolar disorder (BD) patients in the immune marker sample grouped according to recent cannabis use (User) or no recent cannabis use (Non-user).

|  | SCZ patients |  |  |  |  |  | BD patients |  |  |  |  |  |
| --- | --- | --- | --- | --- | --- | --- | --- | --- | --- | --- | --- | --- |
|  | Non-user<br>n=341 |  | User<br>n=60 |  | Test statistic | p-value | Non-user<br>n=205 |  | User<br>n=37 |  | Test statistic | p-value |
|  | Mean | SD | Mean | SD |  |  | Mean | SD | Mean | SD |  |  |
| Age | 31.9 | 10.5 | 26.0 | 7.7 | t=-5.18 | 1.15e-6** | 35.5 | 12.6 | 29.2 | 9.3 | t=-3.63 | 5.67e-4** |
| Sex, female (n, %) | 161 | 46.5 | 16 | 26.7 | $\chi^2=7.42$ | 0.006* | 125 | 61.0 | 18 | 48.6 | $\chi^2=1.49$ | 0.222 |
| Education (years) | 12.9 | 2.9 | 12.0 | 2.1 | t=-2.75 | 0.007* | 14.5 | 3.0 | 13.7 | 3.0 | t=-1.41 | 0.165 |
| BMI | 27.0 | 5.6 | 24.5 | 3.7 | t=-4.17 | 6.54e-5** | 25.6 | 4.5 | 25.4 | 4.7 | t=-0.31 | 0.756 |
| Tobacco use, yes (n, %) | 161 | 46.5 | 52 | 86.7 | $\chi^2=34.66$ | 3.93e-9** | 100 | 48.8 | 26 | 70.3 | $\chi^2=4.97$ | 0.026* |
| Alcohol units 6 months (MD, range) | 10 | 0-1456 | 65 | 0-1098 | W=5769.5 | 6.91e-7** | 52 | 0-1820 | 117 | 2-2184 | W=2404.5 | 0.001* |
| Age of onset (years) | 24.5 | 8.6 | 20.8 | 6.1 | t=-3.96 | 1.35e-4** | 22.8 | 9.9 | 21.4 | 7.7 | t=-1.01 | 0.318 |
| PANSS positive | 15.2 | 5.4 | 17.1 | 5.3 | t=2.60 | 0.011* | 10.2 | 3.9 | 11.7 | 4.1 | t=2.10 | 0.041* |
| PANSS negative | 15.8 | 6.3 | 16.7 | 6.8 | t=0.90 | 0.372 | 10.4 | 3.8 | 9.0 | 2.7 | t=-2.68 | 0.009* |
| PANS general | 31.9 | 9.0 | 34.4 | 8.3 | t=2.04 | 0.044* | 26.1 | 6.8 | 25.2 | 6.2 | t=-0.72 | 0.477 |
| PANSS total | 63.0 | 17.2 | 68.1 | 17.5 | t=2.10 | 0.039* | 46.6 | 11.9 | 45.4 | 10.6 | t=-0.60 | 0.548 |
| IDS | 18.8 | 12.9 | 14.5 | 11.7 | t=-1.88 | 0.068 | 16.1 | 11.5 | 16.0 | 12.0 | t=-0.07 | 0.946 |
| YMRS | 5.1 | 5.0 | 6.3 | 5.6 | t=1.33 | 0.189 | 3.7 | 5.1 | 4.5 | 5.1 | t=0.87 | 0.387 |
| GAF Symptoms | 42.7 | 11.3 | 40.1 | 12.1 | t=-1.54 | 0.127 | 56.6 | 12.5 | 56.2 | 10.1 | t=-0.19 | 0.846 |
| GAF Function | 43.7 | 11.0 | 40.2 | 9.8 | t=-2.48 | 0.015* | 53.2 | 12.7 | 53.0 | 11.8 | t=-0.09 | 0.925 |
| Antipsychotics (DDD) | 1.23 | 1.22 | 1.44 | 0.99 | t=1.47 | 0.146 | 0.53 | 0.76 | 0.34 | 0.50 | t=-1.90 | 0.061 |
| Anticonvulsants (DDD) | 0.084 | 0.26 | 0.057 | 0.24 | t=-0.79 | 0.432 | 0.32 | 0.52 | 0.24 | 0.41 | t=-1.09 | 0.282 |
| Antidepressants (DDD) | 0.47 | 0.85 | 0.28 | 0.55 | t=-2.24 | 0.027* | 0.51 | 0.89 | 0.42 | 0.78 | t=-0.66 | 0.509 |
| Lithium (DDD) | 0.026 | 0.18 | 0.017 | 0.13 | t=-0.48 | 0.630 | 0.24 | 0.50 | 0.20 | 0.46 | t=-0.52 | 0.604 |

\*\*p<0.001, \*p<0.05. BMI: Body Mass Index, PANSS: Positive and Negative Syndrome Scale, IDS: Inventory of Depression Scale, YMRS: Young Mania Rating Scale, GAF: Global Assessment of Functioning, MD: Median, DDD: Defined Daily Dose

### Significant association between cannabis use and sgp130 levels

We first compared the levels of 13 stable plasma biomarkers involved in immune and neuroendocrine regulation (YKL40, CatS, sTNFR1, BDNF, sgp130, IL-1RA, Alcam, MPO, CXCL16, Park7, vWF, OPG, PTX3) in the cannabis user versus non-user groups among BD and SCZ patients. The circulating levels of sgp130 were significantly higher among cannabis users in the SCZ group after adjustment for the number of tests performed (p=0.002; Figure 1, Table 2). The log2 mean difference was 0.175, which translates to an increase of 27.3 ng/mL (12.9%) in absolute concentration. Non-user SCZ patients had a mean sgp130 level below the mean concentration found in healthy controls, but this deficiency was completely reversed in the cannabis group (Figure 1). Females had on average lower sgp130 levels than males, and this difference was generally not affected by cannabis use, indicating that cannabis had similar effects on sgp130 levels in both genders. To further explore the nature of the sgp130 association in SCZ, patients were grouped according to the severity of cannabis use. However, no dose-dependent relationship was found between cannabis use and sgp130 (*F*(2, 42)=0.44, p=0.65), which could be explained by the lack of temporal resolution in the cannabis data (Supplementary Figure S1). Although tobacco smoking was strongly correlated with cannabis use, the duration and frequency of tobacco smoking did not have a significant effect on sgp130 concentration (p=0.75; Supplementary Figure S2). No significant difference in sgp130 levels between users and non-users was observed in the BD group (p=0.87; Figure 1, Table 2). All BD patients, independent of cannabis status, had suboptimal sgp130 levels comparable to the levels found in non-user SCZ patients (Figure 1).

**Figure 1.**
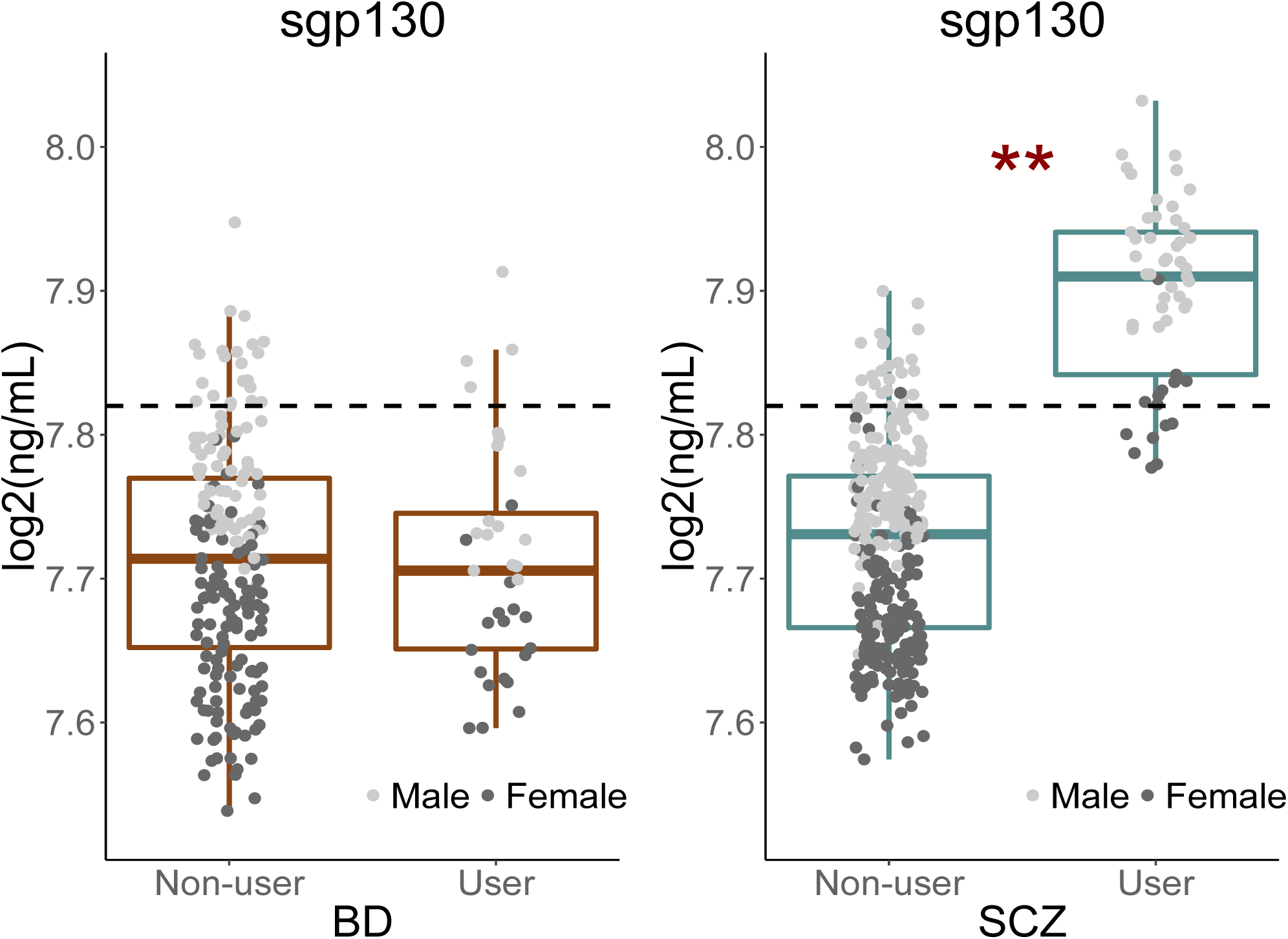
Effect of cannabis use on soluble gp130 levels in bipolar disorder and schizophrenia patients. Cannabis use was significantly associated with increased levels of soluble gp130 in schizophrenia (SCZ) but not in bipolar disorder (BD) patients. The dotted lines represent the mean plasma level of gp130 in healthy controls (n=613). The average gp130 level in BD patients, in both cannabis users and non-users, and in non-user SCZ patients was below the average level found in healthy controls. However, SCZ patients who have used cannabis had an elevated gp130 level above the level for healthy controls. As the dotplots indicate, cannabis use had a comparable up-regulating effect in both males and females. **Bonferroni-adjusted p-value <0.01.

**Table 2.** Associations between inflammatory and neuroendocrine markers in plasma and cannabis use in schizophrenia and bipolar disorder patients.

| Immune marker | SCZ patients |  |  |  |  |  | BD patients |  |  |  |  |  |
| --- | --- | --- | --- | --- | --- | --- | --- | --- | --- | --- | --- | --- |
|  | p-value<br>Nominal | p-value<br>Bonferroni | Non-user |  | User |  | p-value<br>Nominal | p-value<br>Bonferroni | Non-user |  | User |  |
|  |  |  | Mean | SD | Mean | SD |  |  | Mean | SD | Mean | SD |
| sgp130 | 1.62e-4** | 2.11e-3 | 7.724 | 0.067 | 7.899 | 0.063 | 0.866 | 1.000 | 7.712 | 0.081 | 7.712 | 0.079 |
| IL-1RA | 5.91e-3* | 0.077 | 7.783 | 0.424 | 8.241 | 0.304 | 0.255 | 1.000 | 7.711 | 0.500 | 7.285 | 0.572 |
| YKL40 | 6.93e-3* | 0.090 | 5.231 | 0.236 | 5.480 | 0.178 | 0.279 | 1.000 | 5.395 | 0.404 | 5.341 | 0.324 |
| CatS | 0.013* | 0.165 | 2.346 | 0.216 | 2.496 | 0.195 | 0.392 | 1.000 | 2.276 | 0.157 | 2.307 | 0.149 |
| BDNF | 0.020* | 0.255 | 2.230 | 0.174 | 2.534 | 0.142 | 0.259 | 1.000 | 2.254 | 0.181 | 2.072 | 0.197 |
| sTNFR1 | 0.031* | 0.400 | 0.839 | 0.240 | 0.984 | 0.127 | 0.145 | 1.000 | 0.831 | 0.176 | 0.882 | 0.197 |
| Park7 | 0.182 | 1.000 | 2.314 | 0.196 | 2.201 | 0.154 | 0.947 | 1.000 | 2.131 | 0.000 | 2.111 | 0.208 |
| MPO | 0.247 | 1.000 | 8.020 | 0.190 | 8.280 | 0.190 | 0.418 | 1.000 | 8.201 | 0.382 | 7.827 | 0.407 |
| Alcam | 0.327 | 1.000 | 38.619 | 2.694 | 38.205 | 2.104 | 0.756 | 1.000 | 38.880 | 2.224 | 37.160 | 2.454 |
| vWF | 0.379 | 1.000 | 6.335 | 0.226 | 6.214 | 0.167 | 0.352 | 1.000 | 6.126 | 0.308 | 6.356 | 0.348 |
| OPG | 0.400 | 1.000 | 0.420 | 0.112 | 0.418 | 0.085 | 0.082 | 1.000 | 0.485 | 0.148 | 0.305 | 0.156 |
| PTX | 0.500 | 1.000 | 1.576 | 0.250 | 1.833 | 0.205 | 0.600 | 1.000 | 1.530 | 0.238 | 1.476 | 0.222 |
| CXCL16 | 0.701 | 1.000 | 4.060 | 0.162 | 4.099 | 0.150 | 0.059 | 0.769 | 3.972 | 0.156 | 4.156 | 0.121 |

\*\*Significantly associated after Bonferroni correction at $\alpha=0.05$ . \*Nominally associated at $p<0.05$ . The means and standard deviations (SD) shown are for the fitted values from the full regression model.

### Nominally significant cannabis associations

In the SCZ group, nominally significant associations (i.e. associations that did not survive multiple testing correction) were found between cannabis exposure and five inflammatory markers: IL-1RA (p=0.0059), YKL40 (p=0.0069), CatS (p=0.013), sTNFR1 (p=0.031), and the neurotrophin BDNF (p=0.020; Figure 2, Table 2). Although cannabis users had increased levels of all five markers compared to non-users, IL-1RA and BDNF levels were still below the average concentration found in healthy controls. Females had generally higher levels of IL-1RA and lower levels of CatS than males, while no gender differences were seen for YKL40, BDNF, and sTNFR1 (Figure 2).

**Figure 2.**
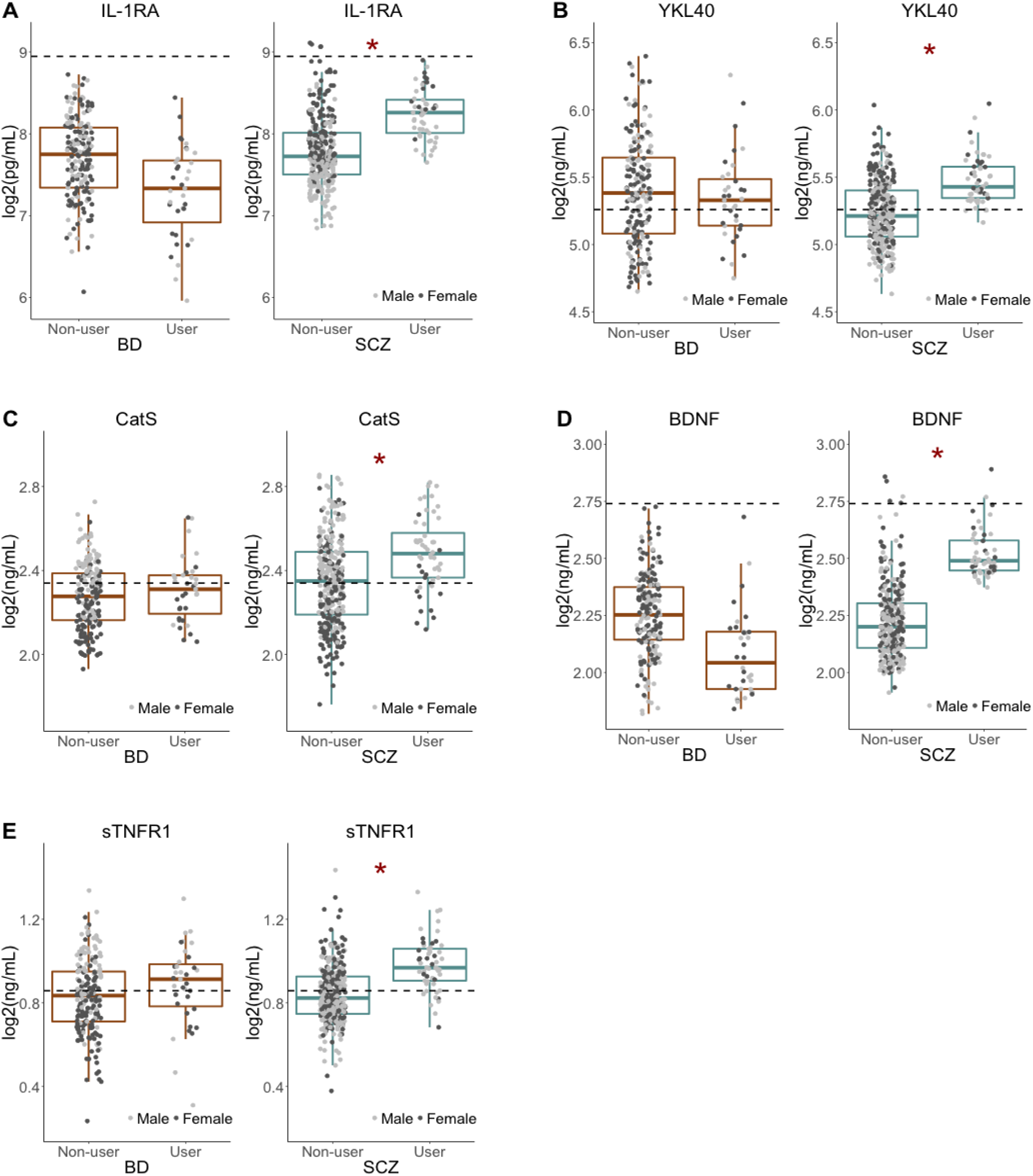
Nominally significant associations between cannabis use and inflammatory and neuroendocrine markers. Cannabis use had a nominally significant association (p<0.5) with five immune and neuroendocrine markers in SCZ but not in BD patients. All markers had higher plasma levels in cannabis users than in non-users. The dotted lines represent the average concentration in healthy controls (n=613). *Nominal p-value <0.05.

### Associations between cannabis use and mRNA expression

To examine if the observed changes in systemic markers could be attributed to the modulation of circulating immune cells by cannabis exposure, we tested whether cannabis status was also associated with the mRNA expression levels of *IL6ST*, *IL1RN, CHI3LI, CTSS, BDNF*, and *TNFRSF1A* genes encoding sgp130, IL-1RA, YKL40, CatS, BDNF, and sTNFR1, respectively. No significant associations were found for any of the genes in BD or SCZ patients (Supplementary Figure S4; Supplementary Table S2).

## DISCUSSION

The aim of the present study was to investigate whether self-administered cannabis use was associated with changes in circulating immune and neuroendocrine markers in SCZ and BD. We performed a neuroimmune screening of 13 plasma markers in SCZ and BD patients, subdividing each group into cannabis user and non-user subgroups. We found that sgp130 concentrations were markedly elevated among cannabis users in the SCZ but not in the BD group. We also found indications of increased levels of IL-1RA, YKL40, CatS, sTNFR1, and BDNF among cannabis using SCZ patients, but these differences were not significant after Bonferroni correction. These results may therefore reflect individual differences in cannabis-related inflammatory functions within and between the sample groups rather than being SCZ or BD-specific biological features. Changes in the plasma concentrations of sgp130, IL-1RA, YKL40, CatS, sTNFR1, and BDNF were not accompanied by corresponding changes at the gene expression level in whole blood samples indicating limited contribution from peripheral immune cell modulation, but could potentially reflect the involvement of other tissues, such as the liver, vascular endothelium, or glial cells.

The critical importance of the regulation of IL-6 trans-signaling through the blood-brain-barrier has been demonstrated in central nervous system inflammation, as well as in neurodegenerative and neuropsychiatric disorders, such as Alzheimer’s and Parkinson’s (reviewed by Banks *et al*., 1995; Rothaug *et al*., 2016). The regulatory effect of gp130 on inflammation is tightly linked to the pleiotropic nature of the IL-6 system in mammals. Gp130, which exists in both a membrane-bound form (gp130) and in a soluble form (sgp130), is a common signal-transducing subunit of several cytokine receptors involved in inflammation and immunity including but not limited to IL-6 (Taga *et al*., 1997). Although IL-6 can be produced by several immune and non-immune cells, its receptor (IL-6R) expression is restricted to only a subset of cells, such as certain lymphocytes (Oberg *et al*., 2006), microglia (Hsu *et al*., 2015), nd hepatocytes (Castell *et al*., 1988). IL-6R also exists in a soluble form (sIL-6R) produced via alternative splicing or limited proteolysis (Rothaug *et al*., 2016). Signaling of the IL-6/IL-6R complex requires the cell membrane-bound form of the co-receptor gp130. Binding of the complex to gp130 initiates downstream signaling targeting JAK2/STAT3, PI3 kinase, ERK, and other intracellular effector pathways (Silver *et al*., 2010). IL-6 can modulate cellular functions in two different ways (Figure 3): In *classical signaling*, IL-6 binds to IL-6R and then the complex triggers the dimerization of membrane-gp130 leading to downstream effects that are pivotal in the regulation of anti-inflammatory signals (Hibi *et al*., 1990). In IL-6 *trans-signaling*, IL-6 is bound by sIL-6R and the complex can activate distant cells expressing membrane-bound gp130 in a paracrine and endocrine manner which is dominantly pro-inflammatory (Rothaug *et al*., 2016). Gp130 signaling is also involved in neuronal and astrocyte differentiation and survival (Nakashima *et al*., 1999), and in the astrocytic control of CNS neurotoxic inflammation (Haroon *et al*., 2011; Sofroniew, 2015).

**Figure 3.**
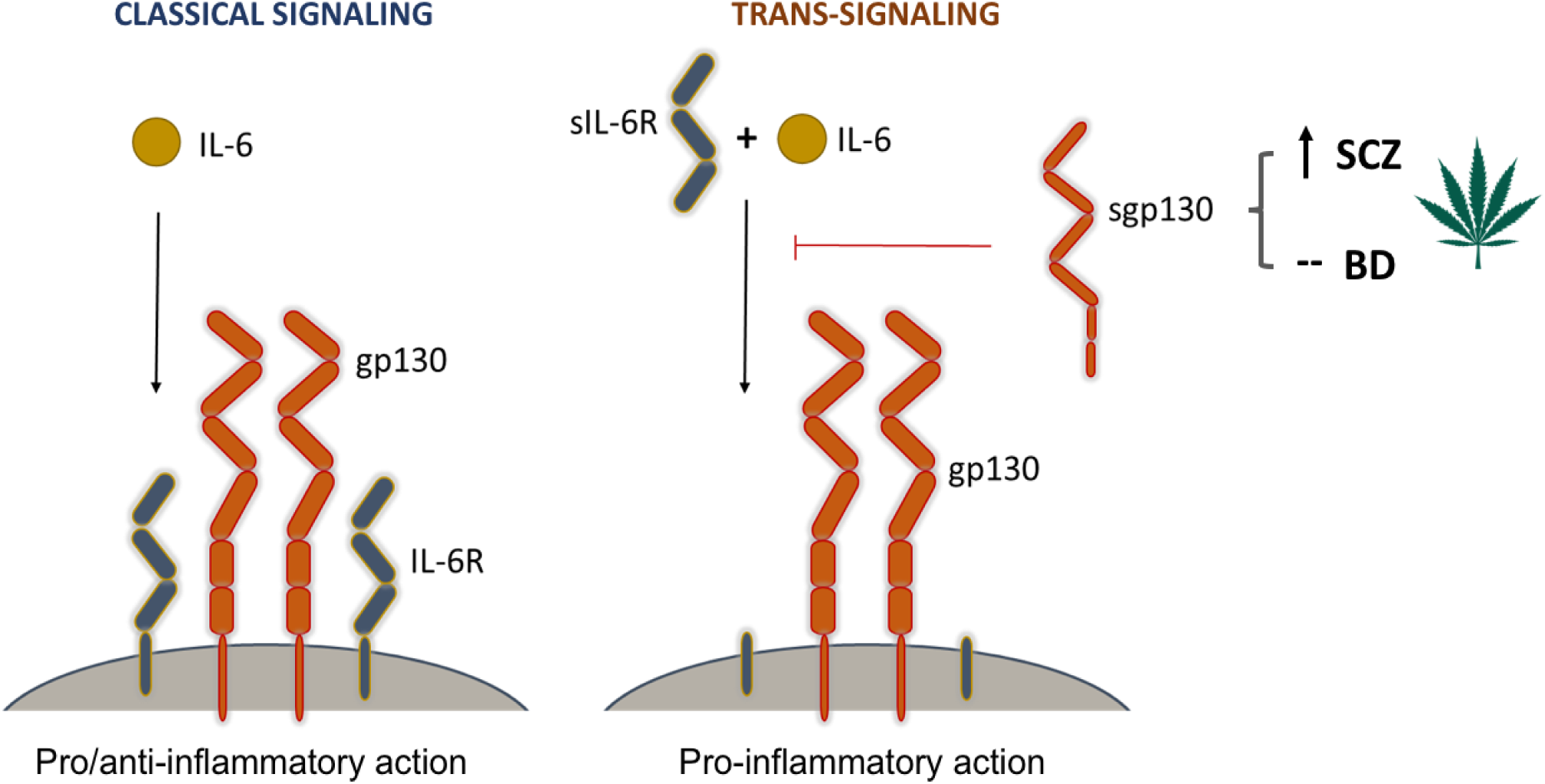
The modulatory effect of cannabis on signaling pathways mediated by gp130. Interleukin-6 (IL-6) is a central player in both pro-inflammatory and anti-inflammatory immune responses. It can modulate cellular responses in two ways. In *classical signaling*, it binds to its cell membrane-bound receptor (IL-6R) and triggers a heterodimeric association with two membrane-bound gp130 molecules. This signaling complex initiates downstream pro- and anti-inflammatory responses through the activation of three signaling cascades, most prominently the JAK-STAT pathway. In *trans-signaling*, IL-6 binds to the soluble form of its receptor (sIL-6R) before it forms a complex with membrane-bound gp130. This complex then activates downstream pathways leading to pro-inflammatory responses. Soluble gp130 (sgp130) has an inhibitory effect on trans-signaling by blocking the association of the IL-6/sIL-6R complex with membrane-bound gp130 molecules. Cannabis use, which was found to increase the levels of sgp130 in schizophrenia patients, is thus believed to have a suppressing effect on inflammation by inhibiting the initiation of trans-signaling cascades.

Soluble gp130 is a decoy receptor that is produced under physiological conditions and is able to bind the IL-6/sIL-6R complex and thereby potently and selectively blocking inflammatory trans-signaling (Narazaki *et al*., 1993; Silver et al., 2010) (Figure 3). Thus, it can alleviate the harmful in vivo effects of IL-6 in the CNS (Campbell *et al*., 2014). Indeed, sgp130 has been shown to exert strong anti-inflammatory effects in both local and systemic contexts (Banks *et al*., 1995; Burton *et al*., 2011; Campbell *et al*., 2014; Rothaug *et al*., 2016), and to protect from the effects of lipopolysaccharide (LPS)-induced sickness behavior in a murine model accompanied by dramatic decrease in IL-6 mRNA and protein levels in the hippocampus (Burton *et al*., 2011).

In accordance with the immune hypothesis of SCZ, dysregulated and/or low-grade chronic brain inflammation may interfere with cognition, mood regulation, and higher-order functioning, and any means of modulating this “inflammatory immune-brain” cross-talk may pose a potential therapeutic target (Khandaker *et al*., 2015; Müller, 2018). In a recent study, sgp130 levels were found to be lower in SCZ and BD patients than in healthy controls (Aas *et al*., 2017). We hypothesize that cannabis consumption may contribute to elevated levels of sgp130 in SCZ patients favoring a systemic anti-inflammatory milieu. This effect may be more pronounced if it is also accompanied by mild increase in the levels of the anti-inflammatory regulators IL-1RA and sTNFR1. Supportive of this idea, a recent report revealed that cannabis use disorder is associated with decreased risk of diseases of gut-brain interaction and inflammatory bowel disease in SCZ, but not in healthy controls (Olesen *et al*., 2019). Since circulating inflammatory biomarker levels are inversely correlated with negative symptom severity in SCZ (Khandaker *et al*., 2015; Müller, 2018), the documented anti-inflammatory effect of cannabis may stand in the background of the observed associations between cannabis consumption and improved mood regulation.

The observed sgp130-modulating effect of cannabis in SCZ may be due to its immunoregulatory potential, especially its effect on IL-6 signaling and presumably on IL-6 trans-signaling feedback-loops as suggested by others (Olah *et al*., 2017). The beneficial effect of cannabis consumption on people with SCZ (“cannabis self-medication hypothesis”) could be attributed to CBD, its natural anti-inflammatory and antipsychotic component, which has recently been reported to be clinically effective in the adjunctive therapy of SCZ (Leweke *et al*., 2012; McGuire *et al*., 2018). Due to its observational nature, our study does not establish a causal link between cannabis use and symptom severity or cognitive performance in SCZ, but may reveal a disease-specific phenomenon that is not present in BD. Additionally, the lack of temporal resolution with regards to our cannabis data (i.e. the lack of control of timing relative to last cannabis use) poses a major limitation to this study. Blood concentrations of the identified immune markers are expected to be affected by both the frequency and amount of cannabis use, as well as the time span that has elapsed since last exposure. The fact that temporal differences were not accounted for could explain why we did not see a dose-dependent relationship between sgp130 levels and cannabis use after stratification by severity of use (use without disabilities, abuse, and addicted). Therefore, although our findings suggest an anti-inflammatory effect of sgp130 that may contribute to the therapeutic value of cannabis in SCZ, elucidation of the physiological details requires future investigations, preferably by comparing sgp130 levels directly with urine concentrations of cannabinoids and their metabolites in SCZ and BD patients.

The absence of the inflammation-modulatory phenomenon of cannabis in BD may resonate with earlier reports on the neutral or potentially adverse effects of cannabis use on BD symptoms and disease evolution (Lagerberg *et al*., 2014; Lagerberg *et al*., 2016). Furthermore, we speculate that our findings regarding the effect of cannabis self-administration on circulating immune marker levels likely reflect the relatively high CBD content of available cannabis products in the streets of Norway (Havig *et al*., 2017). On the other hand, the impact of the psychotropic cannabinoid THC on human brain functions has been highly controversial. It is important to note that, despite its general anti-inflammatory effect, exposure to THC, particularly during adolescence, is hypothesized to perturb the neuroendocrine-immune homeostasis evoking latent vulnerability to psychosis or exacerbate symptoms in individuals with SCZ (Suarez-Pinilla *et al*., 2014; Sherif *et al*., 2016; Di Forti *et al*., 2019). In agreement with this, a recent GWAS study involving a large cohort of lifetime cannabis users revealed causal positive influence between cannabis use and SCZ risk (Pasman *et al*., 2018). Thus, while medical cannabis and cannabinoids are emerging as potential therapeutic adjuvants in neuropsychiatric disorders and other diseases, it is very important to make efforts estimating the possible risks and benefits of cannabinoids in order to provide reliable and competent clinical care.

Taken together, our results show for the first time that cannabis self-administration is associated with elevated levels of sgp130 in SCZ, but not in BD, and that this phenomenon is independent of the modulation of peripheral immune cells. These findings warrant further investigation into the potential neuroimmune, anti-inflammatory, and biobehavioral-cognitive effects of cannabis use in SCZ.

