## Supplementary material for "Cannabis use is associated with increased levels of soluble gp130 in schizophrenia but not in bipolar disorder"

**SUPPLEMENTARY INFORMATION**

SUPPLEMENTARY FIGURES

**Supplementary Figure S1**

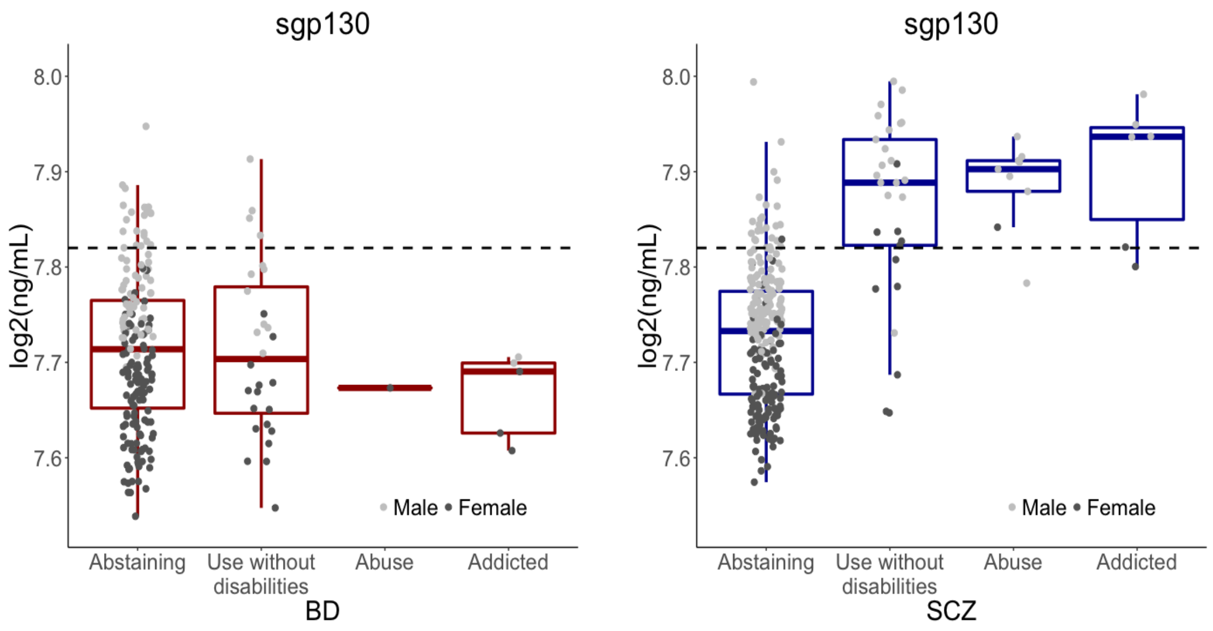


**Figure S1. Association between frequency of cannabis use and sgp130 levels in BD and SCZ patients.** Patterns of cannabis use during the last six months before blood sampling were assessed with the Clinician Drug Use Scale (CDUS). No significant difference in sgp130 concentration was found between the different subgroups of cannabis users in the SCZ group ($F(2,42)=0.44$, $p=0.65$). The lack of a dose-response relationship between cannabis exposure and sgp130 concentration could be due to the lack of temporal resolution in the cannabis data. Since sgp130 levels are expected to depend on the time interval that has passed since the last exposure to cannabis, the inability to account for temporal differences may mask the dose-dependent effect of cannabis on sgp130 concentrations.

**Supplementary Figure S2**

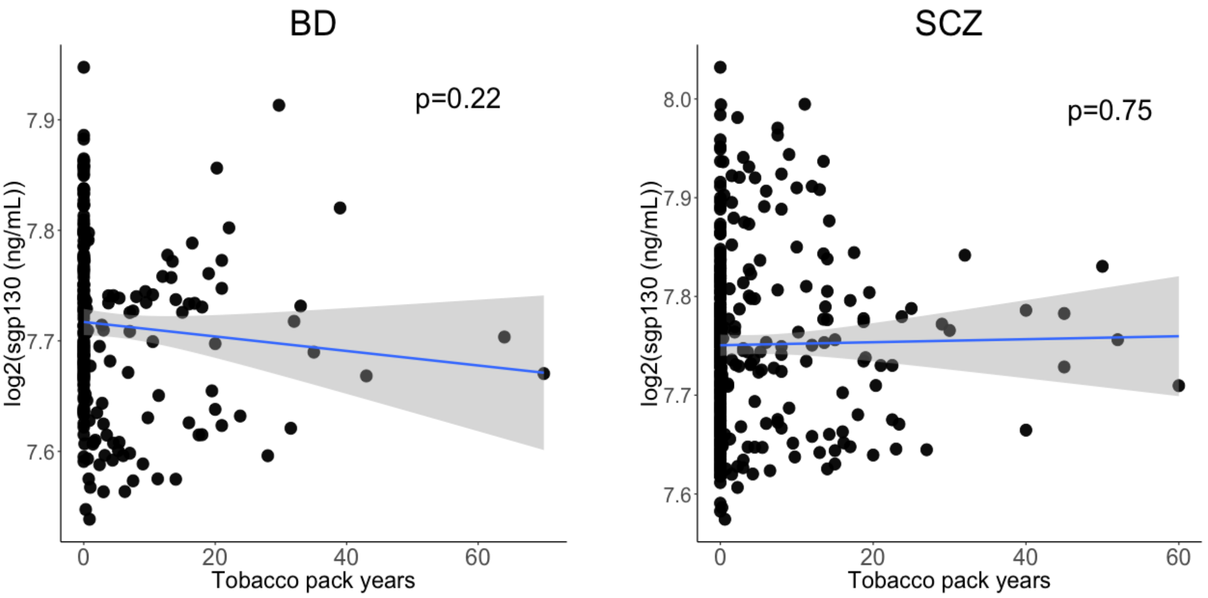


**Figure S2. Relationship between sgp130 and tobacco pack-years.** The figures show the relationship between plasma levels of sgp130 and tobacco pack-years in BD and SCZ patients. Tobacco pack-years is a measure of how much and for how long an individual has been smoking tobacco. Pack-years were defined as the number of cigarettes currently smoked per day multiplied by the number of years the individual has been smoking and divided by 20. No significant relationships were found between sgp130 and tobacco pack-years in BD or SCZ patients.

**Supplementary Figure S3**

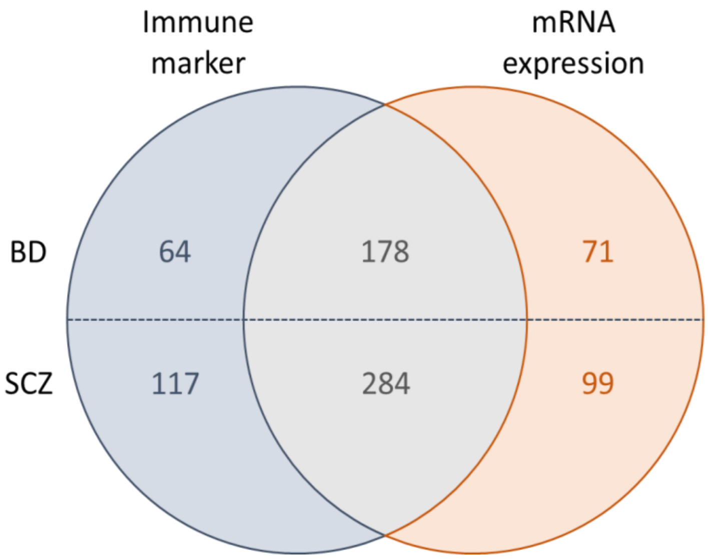


**Figure S3. Partly overlapping immune marker and mRNA expression samples.** To investigate whether the significantly associated serum markers were correlated with cannabis use at the gene expression level, a partly overlapping sample of 249 BD and 383 SCZ patients were used to test for associations between gene expression levels and cannabis exposure. “Immune marker” represents the study sample used to analyze inflammatory markers in serum, while “mRNA expression” represents the study sample used to analyze gene expression levels.

**Supplementary Figure S4**

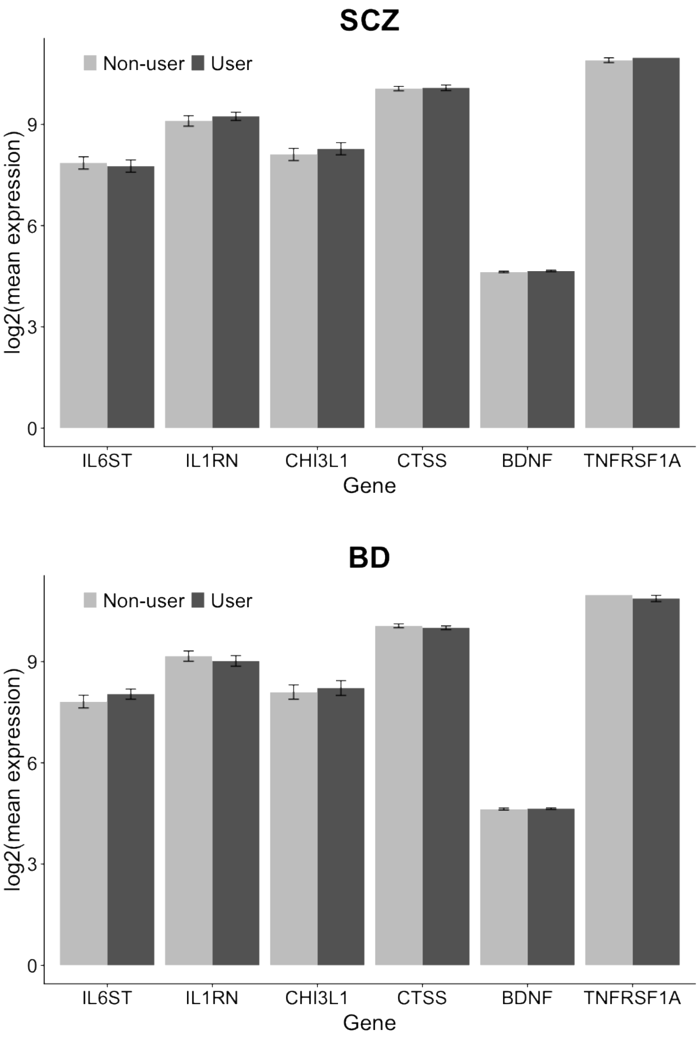


**Figure S4. Effect of cannabis use on gene expression levels of inflammatory markers.** Expression levels of the genes encoding the cannabis-related inflammatory immune markers were tested for associations with cannabis exposure. No significant association was found for any of the six genes in BD or SCZ patients. This finding suggests that the observed effects of cannabis on inflammatory markers do not involve peripheral immune cell modulation, but rather reflect the involvement of other tissues and cell types. The correspondence between gene symbol and immune marker is as follows: *IL6ST* – sgp130, *IL1RN* – IL-1RA, *CHI3L1* – YKL40, *CTS* – CatS, *BDNF* – BDNF, *TNFRSF1A* – sTNFR1.

**Supplementary Table S1. Demographic and clinical characteristics of schizophrenia (SCZ) and bipolar disorder (BD) patients in the mRNA expression sample grouped according to recent cannabis use (User) or no recent cannabis use (Non-user).**

| | **SCZ patients** | | | | | | **BD patients** | | | | | |
|---|---|---|---|---|---|---|---|---|---|---|---|---|
| | Non-user n=321 | | User n=62 | | | | Non-user n=208 | | User n=41 | | | |
| | Mean | SD | Mean | SD | Test statistic | p-value | Mean | SD | Mean | SD | Test statistic | p-value |
| Age | 31.7 | 9.9 | 26.2 | 7.1 | t=5.14 | 1.16e-6** | 35.8 | 12.13 | 28.12 | 8.17 | t=5.02 | 3.08e-6** |
| Sex, female (n, %) | 142 | 44.2 | 20 | 32.3 | $\chi^2$=2.58 | 0.108 | 128 | 61.5 | 17 | 41.5 | $\chi^2$=4.88 | 0.027* |
| Education (years) | 13.1 | 2.9 | 12.3 | 2.2 | t=2.42 | 0.017* | 14.6 | 3.1 | 13.7 | 2.5 | t=1.99 | 0.051 |
| BMI | 26.9 | 5.6 | 25.1 | 4.4 | t=2.76 | 0.007* | 25.8 | 4.4 | 25.5 | 4.7 | t=0.40 | 0.688 |
| Tobacco use, yes (n, %) | 155 | 48.3 | 51 | 82.3 | $\chi^2$=23.69 | 1.13e-6** | 109 | 52.4 | 29 | 70.7 | $\chi^2$=3.94 | 0.047* |
| Alcohol units 6 months (MD, range) | 13 | 0-1456 | 40 | 0-1098 | W=6561.5 | 2.11e-4** | 37.5 | 0-1820 | 143 | 2-2184 | W=2555 | 1.10e-4** |
| Age of onset (years) | 24.7 | 8.6 | 20.3 | 5.6 | t=5.12 | 1.14e-6** | 23.2 | 9.7 | 20.8 | 7.0 | t=1.89 | 0.063 |
| PANSS positive | 15.1 | 5.3 | 16.4 | 5.2 | t=-1.87 | 0.066 | 10.0 | 3.8 | 10.8 | 3.9 | t=-1.21 | 0.230 |
| PANSS negative | 15.8 | 5.8 | 15.6 | 6.2 | t=0.20 | 0.839 | 10.4 | 3.7 | 9.6 | 3.7 | t=1.20 | 0.236 |
| PANS general | 31.8 | 8.5 | 33.7 | 8.3 | t=-1.66 | 0.099 | 26.0 | 6.5 | 24.9 | 5.7 | t=1.14 | 0.261 |
| PANSS total | 62.7 | 15.9 | 65.8 | 17.1 | t=-1.33 | 0.187 | 46.3 | 11.5 | 45.3 | 10.4 | t=0.59 | 0.559 |
| IDS | 17.2 | 12.3 | 15.1 | 11.8 | t=0.99 | 0.323 | 16.6 | 12.2 | 15.7 | 12.1 | t=0.38 | 0.703 |
| YMRS | 4.8 | 4.8 | 5.5 | 5.5 | t=-0.84 | 0.406 | 3.7 | 5.2 | 3.6 | 4.6 | t=0.16 | 0.870 |
| GAF Symptoms | 43.1 | 11.4 | 40.4 | 11.4 | t=1.68 | 0.097 | 56.5 | 12.0 | 57.4 | 10.4 | t=-0.49 | 0.625 |
| GAF Function | 43.8 | 11.1 | 41.2 | 9.6 | t=1.91 | 0.059 | 53.4 | 12.9 | 54.0 | 12.0 | t=-0.27 | 0.787 |
| Antipsychotics (DDD) | 1.57 | 3.21 | 1.84 | 2.80 | t=-0.68 | 0.500 | 0.69 | 1.56 | 0.50 | 0.83 | t=1.09 | 0.277 |
| Anticonvulsants (DDD) | 0.088 | 0.27 | 0.056 | 0.21 | t=1.04 | 0.303 | 0.32 | 0.50 | 0.26 | 0.40 | t=0.83 | 0.411 |
| Antidepressants (DDD) | 0.45 | 0.87 | 0.38 | 0.65 | t=0.74 | 0.462 | 0.57 | 0.94 | 0.34 | 0.77 | t=1.63 | 0.109 |
| Lithium (DDD) | 0.026 | 0.17 | 0.048 | 0.28 | t=-0.62 | 0.540 | 0.21 | 0.46 | 0.21 | 0.47 | t=-0.03 | 0.973 |

**p<0.001, *p<0.05. BMI: Body Mass Index, PANSS: Positive and Negative Syndrome Scale, IDS: Inventory of Depression Scale, YMRS: Young Mania Rating Scale, GAF: Global Assessment of Functioning, MD: Median, DDD: Defined Daily Dose.

**Supplementary Table S2. Associations between mRNA expression levels of significant serum markers and cannabis use in schizophrenia and bipolar disorder patients.**

| | | SCZ patients | | | | | | BD patients | | | | | |
|---|---|---|---|---|---|---|---|---|---|---|---|---|---|
| | | | | Non-user | | User | | | | Non-user | | User | |
| Gene | Immune marker | p-value Nominal | p-value Bonferroni | Mean | SD | Mean | SD | p-value Nominal | p-value Bonferroni | Mean | SD | Mean | SD |
| *IL6ST* | gp130 | 0.198 | 1.000 | 7.858 | 0.182 | 7.764 | 0.181 | 0.208 | 1.000 | 7.816 | 0.189 | 8.038 | 0.152 |
| *IL1RN* | IL-1RA | 0.091 | 0.546 | 9.103 | 0.156 | 9.240 | 0.123 | 0.781 | 1.000 | 9.163 | 0.153 | 9.021 | 0.158 |
| *CHI3L1* | YKL40 | 0.419 | 1.000 | 8.110 | 0.182 | 8.278 | 0.180 | 0.494 | 1.000 | 8.097 | 0.212 | 8.217 | 0.220 |
| *CTSS* | CatS | 0.721 | 1.000 | 10.063 | 0.066 | 10.085 | 0.082 | 0.711 | 1.000 | 10.060 | 0.057 | 10.004 | 0.055 |
| *BDNF* | BDNF | 0.168 | 1.000 | 4.629 | 0.022 | 4.659 | 0.025 | 0.476 | 1.000 | 4.632 | 0.032 | 4.643 | 0.023 |
| *TNFRSF1A* | sTNFR1 | 0.161 | 0.966 | 10.902 | 0.070 | 10.975 | 0.066 | 0.379 | 1.000 | 10.973 | 0.101 | 10.871 | 0.092 |

The means and standard deviations (SD) shown are for the fitted values from the full regression model.
